# Knocking out histidine ammonia-lyase by using CRISPR-Cas9 abolishes histidine role in the bioenergetics and the life cycle of *Trypanosoma cruzi*

**DOI:** 10.1101/2025.02.19.639176

**Authors:** Janaína de Freitas Nascimento, María Julia Barisón, Gabriela Torres Montanaro, Letícia Marchese, Rodolpho Ornitz Oliveira Souza, Letícia Sophia Silva, Alessandra Aparecida Guarnieri, Ariel Mariano Silber

## Abstract

*Trypanosoma cruzi,* the causing agent of Chagas disease, is the only known trypanosomatid pathogenic to humans having a complete histidine to glutamate pathway, which involves a series of four enzymatic reactions that convert histidine into downstream metabolites, including urocanate, 4-imidazolone-5-propionate, N-formimino-L-glutamate and L-glutamate. Recent studies have highlighted the importance of this pathway in ATP production, redox balance, and the maintenance of cellular homeostasis in *T. cruzi*. In this work, we focus on the first step of the histidine degradation pathway, which is performed by the enzyme histidine ammonia lyase. Here we determined the kinetic and biochemical parameters of the *T. cruzi* histidine ammonia-lyase. By generating null mutants of this enzyme using CRISPR-Cas9 we observed that disruption of the first step of the histidine degradation pathway completely abolishes the capability of this parasite to metabolise histidine, compromising the use of this amino acid as an energy and carbon source. Additionally, we showed that the knockout of the histidine ammonia lyase affects metacyclogenesis when histidine is the only metabolizable source and diminishes trypomastigote infection *in vitro*.

## Introduction

*Trypanosoma cruzi* is the causative agent of Chagas disease, which is a neglected tropical disease and affects millions of people, predominantly in the Americas (WHO, 2023). Despite first described over a century ago by Dr Carlos Chagas (Chagas, 1909), the biology of *T. cruzi* remains incompletely understood. The life cycle of *T. cruzi* involves both a triatomine vector and a vertebrate host. When the triatomine insect feeds on the blood of an infected mammalian host, it ingests non-replicative trypomastigotes. These transform into replicative epimastigotes within the insect’s digestive tract. The epimastigotes colonize the gut, migrates to its posterior section, adhere to the epithelium, and differentiate into the non-replicative metacyclic trypomastigotes. During a subsequent blood meal, the infected triatomine releases metacyclic trypomastigotes in its faeces, which can enter the mammalian host through the bite wound inflicted by the insect or through mucous membranes. Once inside the host, the trypomastigotes invade the mammalian cells and differentiate into the replicative amastigotes. These amastigotes replicate, differentiate back into trypomastigotes, and are either released into the bloodstream or infect neighbouring cells upon host-cell rupture (reviewed by Martín-Escolano et al., 2022).

The metabolism of *T. cruzi* relies on the catabolism of carbohydrates, amino acids, and lipids as carbon and energy sources (Booth & Smith, 2020; Cazzulo, 1992; Marchese et al., 2018; Souza et al., 2021; Urbina, 1994) When glucose is available in the extracellular environment, it is primarily metabolized through glycolysis (Cazzulo, 1994). However, when extracellular glucose levels decrease, amino acids become the main energy source fuelling oxidative pathways that feed electrons into the respiratory chain (Barisón et al., 2016; Mantilla et al., 2015; L. M. Marchese et al., 2020; Paes et al., 2013; Sylvester et al., 1976; Zeledon, 1960). Beyond their role in translation and metabolism, amino acids are involved in a variety of critical biological processes in *T. cruzi*. These include osmotic regulation (Rohloff et al., 2003), protection against apoptosis (Piacenza et al., 2001), regulation of cell differentiation and invasion (Contreras et al., 1985; Krassner et al., 1990; Mantilla et al., 2015; Martins et al., 2009), and resistance to oxidative imbalance and thermal and nutritional stresses (Arias et al., 2011; Magdaleno et al., 2011; Mantilla et al., 2021; Paes et al., 2013).

*T. cruzi* does not synthesize histidine (His) and must obtain it via transport from the extracellular environment. Once internalized, His can be fully oxidized to CO□, potentially supplying electrons to the respiratory chain and promoting ATP synthesis through oxidative phosphorylation (Barisón et al., 2016). Another role that has been attributed to His is the production of ovothiol, a thiolated version of the amino acid which works as a redox defence (Ariyanayagam, 2001; Braunshausen & Seebeck, 2011). Two distinct His catabolic pathways have been described, differentiated by their end products. Both pathways generate ammonia and glutamate (Glu); however, type I pathway additionally produces formamide, while the type II pathway yields formate (Bender, 2012). Among trypanosomatids pathogenic to humans, *T. cruzi* is uniquely characterized by its possession of sequences encoding all four canonical enzymes involved in the His degradation pathway. Specifically, *T. cruzi* encodes enzymes for the type 1 pathway, which is highly conserved in bacteria (Bender, 2012). This pathway begins with the removal of the α-amino group of His by Histidine ammonia-lyase (*Tc*HAL, EC 4.3.1.3), resulting in the production of urocanate and ammonia. Urocanate is subsequently converted into 4-imidazolone-5-propionate (IPA) by urocanate hydratase (*Tc*UH, EC 4.2.1.49) (Retey, 1994). The amide bond of IPA is then hydrolyzed by imidazolonepropionase (*Tc*IP, EC 3.5.2.7), yielding N-formimino-L- glutamate. Finally, the formimino group can be hydrolyzed to produce formamide and Glu via formiminoglutamase (*Tc*FG, EC 3.5.3.8) (Figure 1) (Magasanik & Bowser, 1955). The enzymatic activities of *Tc*HAL, *Tc*UH and *Tc*FG have been experimentally confirmed, and their crystal structures have been solved at high resolution (Boreiko et al., 2020; Hai et al., 2013; Miranda et al., 2020).

**Figure 1.**
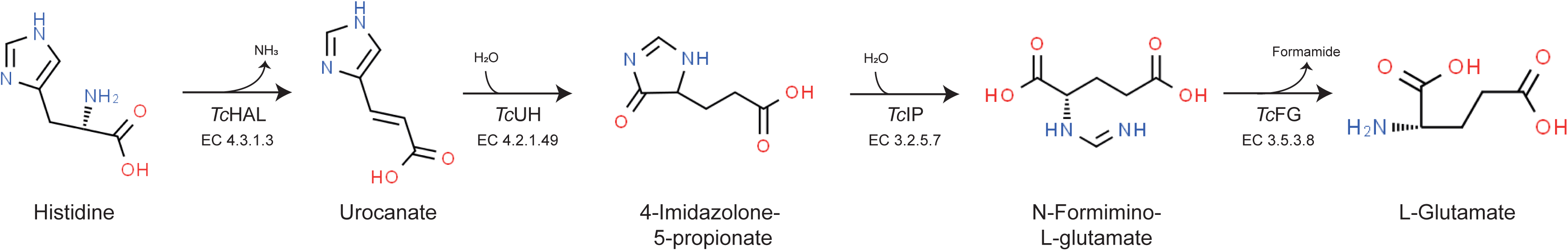
Histidine degradation pathway. In *T. cruzi*, the histidine degradation pathway occurs linearly with no contribution of other metabolic pathways, producing glutamate (Glu) and formamide as end products. The enzymes involved in this pathway include: histidine ammonia-lyase (*Tc*HAL, EC 4.3.1.3), urocanate hydratase (*Tc*UH, EC 4.2.1.49), imidazolonepropionase (*Tc*IP, EC 3.5.2.7), formiminoglutamase (*Tc*FG, EC 3.5.3.8).

The roles of *Tc*HAL the first enzyme of the His degradation pathway in *T. cruzi*, extend beyond His bioenergetic functions related to His metabolism. Recent studies have revealed that *Tc*HAL localizes to both, the cytoplasm and acidocalcisomes in epimastigotes. In these organelles, His deamination contributes to the regulation of intraorganellar pH, a process modulated by polyphosphate (polyP) binding. This function in acidocalcisome alkalinization has been proposed as critical for the survival of epimastigotes under conditions of nutritional stress (Mantilla et al., 2021). Despite the apparent importance of *Tc*HAL in *T. cruzi* biology, its expression and functional impact across the parasite’s life cycle remain unexplored. In this study, we generated *Tc*HAL null mutants, to assess their ability to metabolize His, and to investigate this enzyme’s role in the completion of the parasite’s life cycle.

## Results

### Kinetic and biochemical characterization of recombinant *Tc*HAL

To determine the parameters that govern *Tc*HAL enzymatic activity, the enzyme was heterologously expressed in *Escherichia coli* and purified by affinity chromatography (Supp. Figure 1). The recombinant enzyme was then used to measure its activity over time, with varying concentrations of His as the substrate (Figure 2A). Analysis of the initial velocities (*V*_0_) as a function of substrate concentration revealed that *Tc*HAL follows product formation kinetics consistent with a hyperbolic curve, fitting in the Michaelis-Menten model (Figure 2B). The estimated values for *K*_m_ and *V*_max_ for the recombinant enzyme were 0.77 ± 0.330 mM and 4.03 ± 1.87 μmol·min^−1^·mg^−1^, respectively. When same parameters were measured in epimastigote cell-free extracts, the obtained apparent *K*_m_ and *V*_max_ were 0.32 ± 0.08 mM, and 0.03 ± 0.01 μmol·min^−1^·mg^−1^, respectively (Figure 2B, inset).

**Figure 2.**
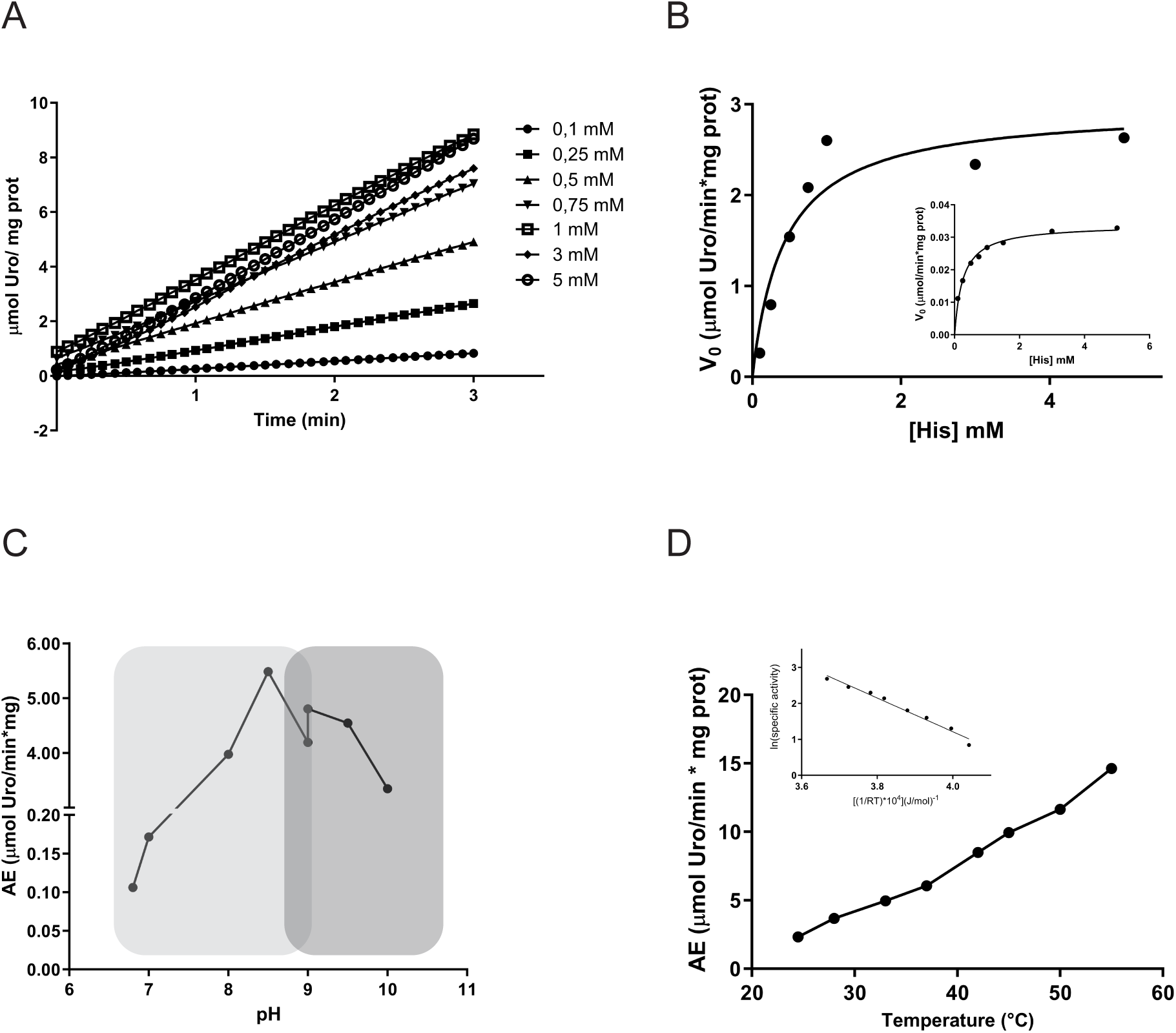
Kinetic parameters of *Tc*HAL. **A.** *Tc*HAL activity as a function of time. Enzyme activity measurements were performed using real-time spectrophotometric assays, monitoring the increase in absorbance at 277 nm, corresponding to the conversion of histidine into the product urocanate. The reaction mixture for *Tc*HAL contained 100 mM Tris-HCl buffer (pH 9), 0.1 mM MnCl□, 1.7 mM reduced glutathione (GSH) and the indicated concentrations of histidine. The reaction was initiated by adding purified recombinant *Tc*HAL enzyme (10 – 50 μg) or epimastigote protein extract (50 – 100 μg). The initial reaction velocity (*V*□) was calculated in the linear region of the reaction curve using the molar extinction coefficient of urocanate (CEM_277nm_ = 1.88 M□¹ cm□¹). **B.** Michaelis-Menten curve for recombinant *Tc*HAL and for the epimastigote protein extract (inset). **C.** pH dependence of recombinant *Tc*HAL activity. Tris-HCl (pH 6.8–9, light grey shading) and CHES (N-cyclohexyl-2- aminoethanesulfonic acid, pH 9–10, dark grey shading) were used as reaction buffers. **D.** Temperature dependence of recombinant *Tc*HAL activity. Enzymatic activity was measured in a temperature range of 25–55 °C. The activation energy value was determined from the Arrhenius plot (inset). Graphs are representative of at least three independent replicates.

Following the kinetic characterization, additional biochemical properties of recombinant *Tc*HAL were assessed, including its dependence on temperature and pH. The results revealed that *Tc*HAL exhibits maximal activity within a basic pH range of 8 to 9 (Figure 2C). Additionally, its activity increased near linearly with temperature between 25°C and 55°C (Figure 2D). Using these data, the activation energy (*E*_a_) was calculated using the Arrhenius equation, yielding a value of 39.8 ± 6.1 kJ·mol^−1^ (Figure 2D, inset).

### Generation and confirmation of *Tc*HAL^-/-^ epimastigotes and add-backs

To investigate the biological role of the His degradation pathway in *T. cruzi*, a knockout cell line for the *Tc*HAL coding sequence was generated using CRISPR-Cas9 technology to disrupts the pathway’s first enzymatic step. The system used herein relies on the constitutive expression of the endonuclease *Sp*Cas9 and the viral RNA polymerase T7 Pol from the tubulin loci in epimastigotes of the CL Brener strain. T7 Pol enables the *in vivo* transcription of the single guide RNA (sgRNA), which is delivered to the parasite as a double-stranded DNA (dsDNA), alongside a recombination cassette containing the desired modification (Costa et al., 2018). For obtaining the *Tc*HAL knockout, two sgRNA sequences were designed: one to target Cas9 cleavage at the start (position +33) and another targeting the end (position +1558) of the *Tc*HAL coding region. A recombination cassette containing the blasticidin resistance gene flanked by 30 bp homologous sequences of the *Tc*HAL coding region, was used to delete the gene of interest (Figure 3A). Following transfection, selection, and cloning, three independent clones with confirmed *Tc*HAL coding sequence deletion were obtained. The deletion was verified by: (i) PCR using *Tc*HAL-specific primers, which showed no amplification of the target band (Figure 3C); and (ii) enzymatic activity assays, which demonstrated the absence of urocanate (Uro), the product of His degradation by *Tc*HAL, in cell-free extracts incubated with His (Figure 3D). To generate add-back cell lines, clone 1 of the *Tc*HAL^-/-^ cell line was randomly selected and transfected with the plasmid *Tc*HAL_pTEX_puroR (Figure 3B). Restoration of *Tc*HAL expression was also confirmed by genotyping and enzymatic activity assays (Figures 3C and 3D).

**Figure 3.**
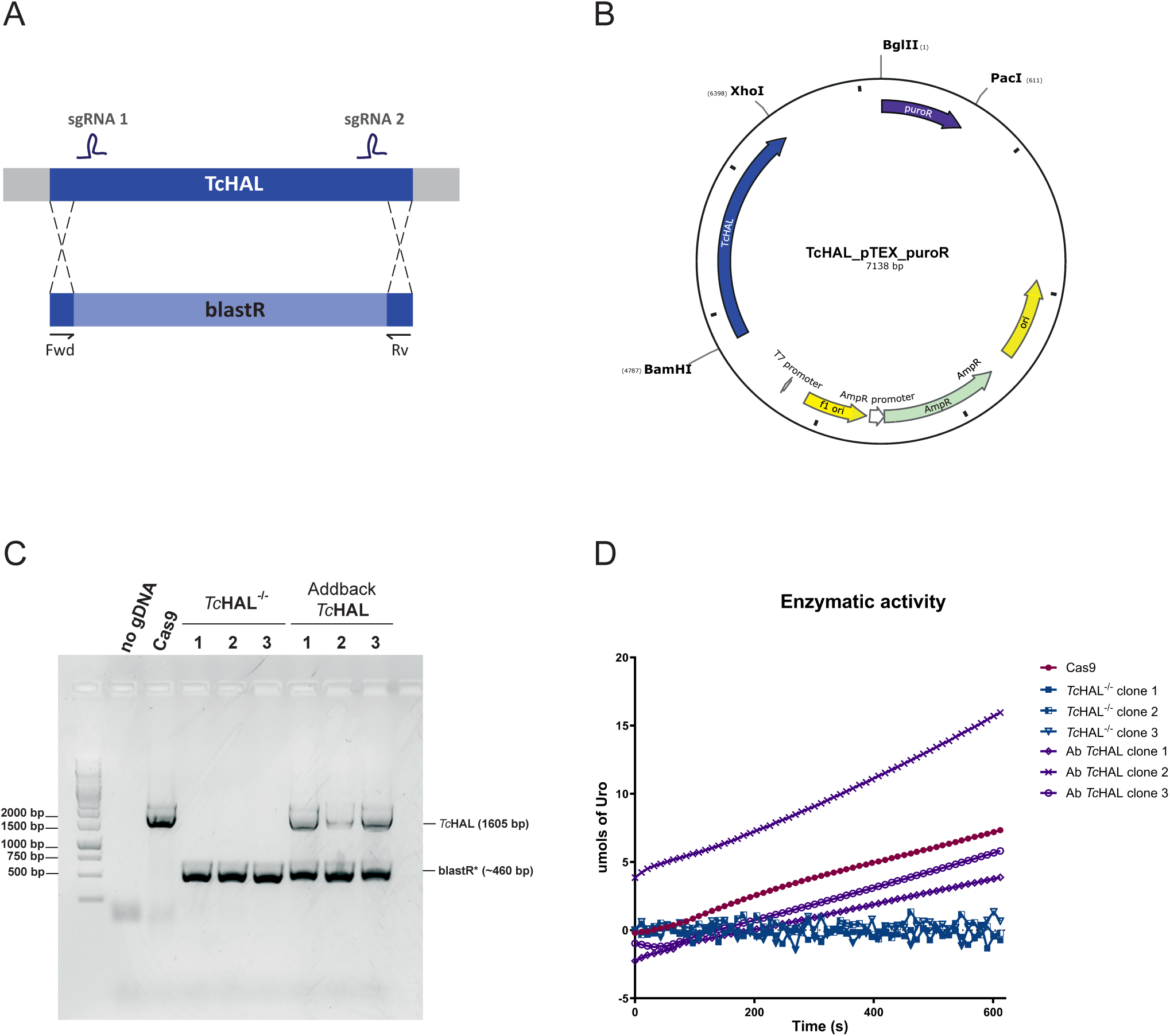
Generation and confirmation of *Tc*HAL^-/-^ epimastigotes using CRISPR- Cas9. **A.** Schematic representation of the CRISPR-Cas9 strategy used to substitute the *Tc*HAL coding sequence (CDS) by the blasticidin resistance gene (blastR). Two single guide RNAs (sgRNAs) were used to direct the Cas9 endonuclease to positions +33 and +1558 in the *Tc*HAL CDS. Primers Fwd and Rev were used for genotyping. **B.** Map of the plasmid *Tc*HAL_pTEX_puroR, which allows episomal expression of *Tc*HAL and contains the puromycin resistance gene as a selectable marker. **C.** Genotyping of three independent clones of the *Tc*HAL^-/-^ and add-back cell lines by PCR. Amplification corresponding to *Tc*HAL CDS (1605 bp) is absent in *Tc*HAL^-/-^ clones, where only the amplification of blastR added of the 60 bp remaining of *Tc*HAL CDS (460 bp) was observed. **D.** Graph shows *Tc*HAL enzymatic activity assay, performed by monitoring the formation of urocanate. *Tc*HAL^-/-^ epimastigotes show no detectable activity, which is restored in the add-back clones.

### *Tc*HAL^-/-^ epimastigotes are not able to use His as an ATP source

As previously mentioned, His is an oxidizable amino acid whose degradation produces intermediates of the tricarboxylic acids cycle (TCAc), which serves as substrates for the respiratory chain. This process drives O_2_ consumption and mitochondrial ATP formation through the oxidative phosphorylation (OxPhos) (Barisón et al., 2016). To investigate whether *Tc*HAL^-/-^ parasites retain the ability to fully oxidize His, a CO_2_ trap assay was performed. Parasites were incubated in the presence of ^14^C (L-[^14^C(1,2)]-His), and ^14^CO_2_ accumulation was measured over time. The results show significant ^14^CO_2_ accumulation in the control lineage (Cas9), which was drastically reduced in *Tc*HAL^-/-^ epimastigotes. This indicates that ablation of the first step of the His degradation pathway by deletion of the *Tc*HAL coding sequence abolishes the parasite’s ability to oxidize His to CO_2_ (Figure 4A). To assess whether the incorporation of His and its degradation products into macromolecules was affected in *Tc*HAL^-/-^ epimastigotes, parasites incubated in the presence of ^14^C (L-[^14^C(1,2)]-His) were fractionated. The supernatant from fractionation contained free metabolites, while macromolecules were precipitated by perchloric acid. Compared to control, *Tc*HAL^-/-^ parasites exhibited a ∼65% reduction in the incorporation of His-derived ^14^C into macromolecules after four hours of incubation. Partial recovery was observed in the add-back cell line, while no differences were detected in the soluble fraction (Figures 4B and 4C).

**Figure 4.**
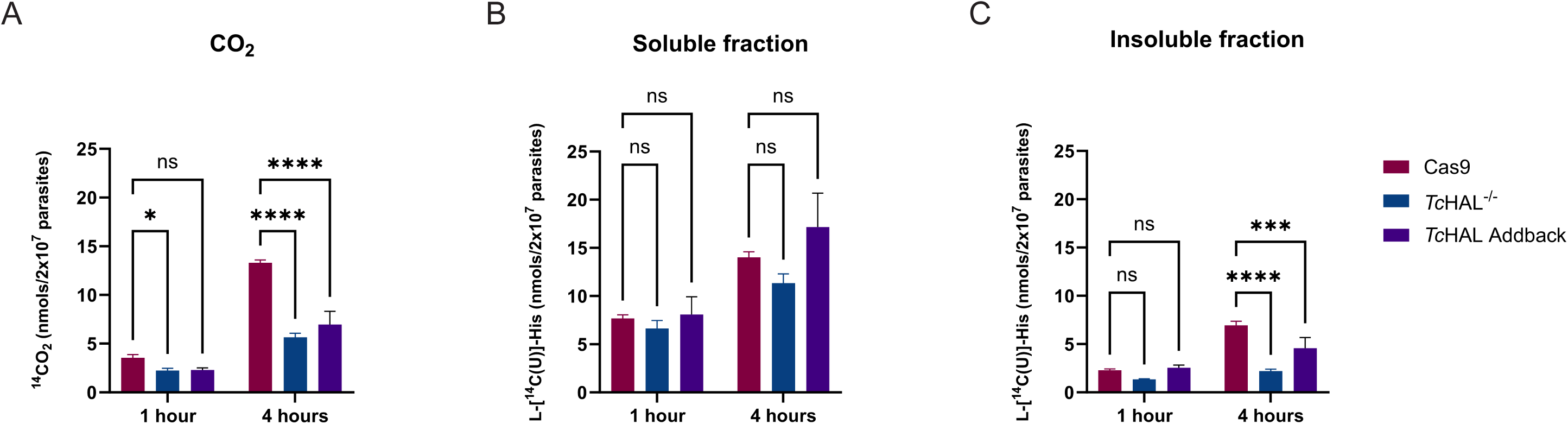
The deletion of *Tc*HAL abolishes the parasites’ ability to oxidize His. **A.** CO□ levels produced from the complete oxidation of His measured by scintillation in Cas9 and *Tc*HAL^-/-^ epimastigotes after incubation with (L-[^14^C(U)]-His). *Tc*HAL^-/-^ parasites show reduced production of CO_2_ from labelled His. **B.** and **C.** Levels of L- [^14^C(1,2)]-His and its degradation products in free metabolites (soluble fraction) and those incorporated into macromolecules (insoluble fraction) measured by scintillation after cell fractionation. *Tc*HAL^-/-^ parasites show reduced incorporation of labelled His and its products into macromolecules. Graphs show the average from experiments performed with 3 independent clones of each cell line. Error bars represent the standard deviation among biological replicates. Statistical analysis was performed using two-way ANOVA with multiple comparisons by Sidák’s test (α=0.05, ns = statistically non-significant difference, * p ≤ 0.05, **** p ≤ 0.0001).

Next, to investigate if inactivation of the first step of the His degradation pathway affects the mutant parasites’ ability to use His as an electron donor for the electron transfer chain (ETC), the respiratory capacity of the parasites was measured following nutritional stress and recovery in the presence of an energy source. Similar to His, proline (Pro) can feed the ETC via its conversion to Glu (Mantilla et al., 2015; Paes et al., 2013). As expected, the use of Pro to trigger cell respiration was not affected by *Tc*HAL knockout (Figure 5A). However, compared to the parental cell line, *Tc*HAL^-/-^ epimastigotes were unable to use His to trigger O_2_ consumption and, consequently, could not produce ATP through the oxidation of this metabolite (Figure 5B). In contrast, *Tc*HAL^-/-^ parasites retained the ability to utilize urocanate, the product of the reaction catalysed by *Tc*HAL, for O_2_ consumption, similar to control lineages (Figure 5C). These results confirm that downstream steps of the His degradation pathway remain functional. When basal respiration and maximum respiratory capacity via ETC were quantified, *Tc*HAL^-/-^ parasites recovered in His exhibited O_2_ flux values comparable to those of the same lineage recovered in respiration buffer without an ATP source (Figure 5D-F).

**Figure 5.**
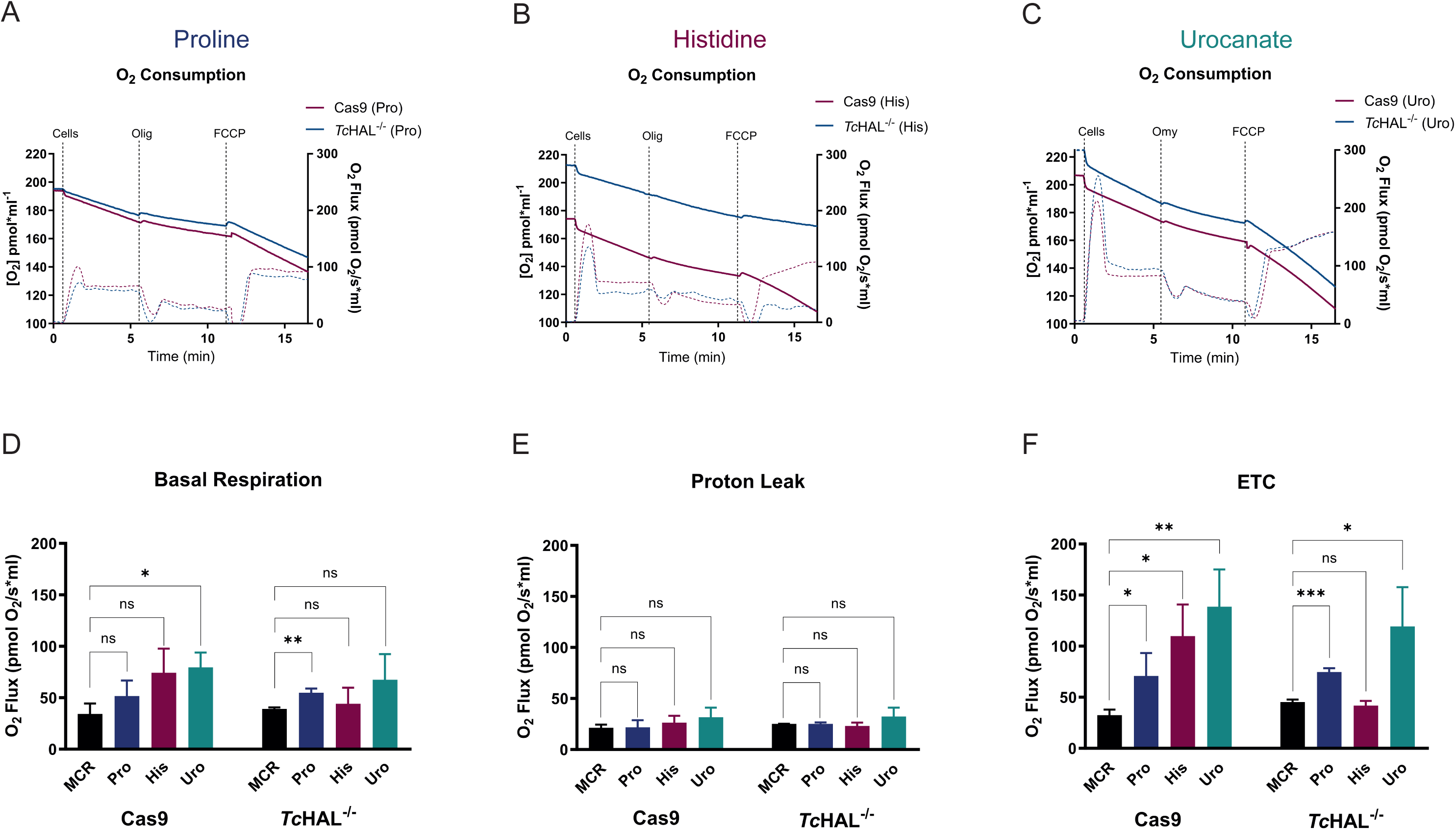
Epimastigotes that do not express *Tc*HAL are unable to use His to trigger O□ consumption. Epimastigotes were pre-stressed in PBS for 16 hours and then recovered for 30 minutes in **A.** Proline (positive control), **B.** Histidine, and **C.** Urocanate. Maximum respiratory capacity was measured following ATP synthase inhibition by oligomycin (Omy) addition and subsequent dissipation of the proton gradient by addition of carbonyl cyanide-p-trifluoromethoxyphenylhydrazone (FCCP). Graphs are representative of one of the three biological replicates performed for each cell line (Cas9 and *Tc*HAL*^-/-^*) in each condition. Quantification of **D.** Basal Respiration, **E.** Proton Leak and **F.** Electron Transport Chain (ETC) shows that *Tc*HAL*^-/-^* epimastigotes recovered in the presence of His have are unable to trigger cell respiration, similarly to parasites recovered only in the MCR buffer. Error bars represent the standard deviation of experiments performed with three independent clones for each strain (Cas9 and *Tc*HAL^-/-^) for each condition. Note the difference in the axis of the graph corresponding to the ETC. Statistical analysis was performed using unpaired t- test, using MCR as control (ns = statistically non-significant difference, * p ≤ 0.05, ** p ≤ 0.01, *** p ≤ 0.001).

Given that His can serve as an important ATP source in *T. cruzi*, we investigated whether epimastigotes lacking *Tc*HAL expression could still utilize His to maintain cell viability during prolonged periods of nutritional stress. Parasites were incubated in PBS supplemented or not with different energy sources (Glc, His, and Uro). Cell viability was assessed using resazurin, a dye that detects cellular redox activity (Ansar Ahmed et al., 1994). While parasites incubated in non-supplemented PBS gradually lost viability over time, those maintained in PBS supplemented with Glc, His or Uro remained viable for up to 72 hours (Figure 6A-C). However, *Tc*HAL^-/-^ parasites maintained in PBS supplemented with His lost viability at a rate similar to those in PBS alone, indicating their inability utilize His to survive nutritional stress (Figure 6B). This phenotype was fully restored in the add-back cell line, where *Tc*HAL expression was reintroduced (Figure 6C).

**Figure 6.**
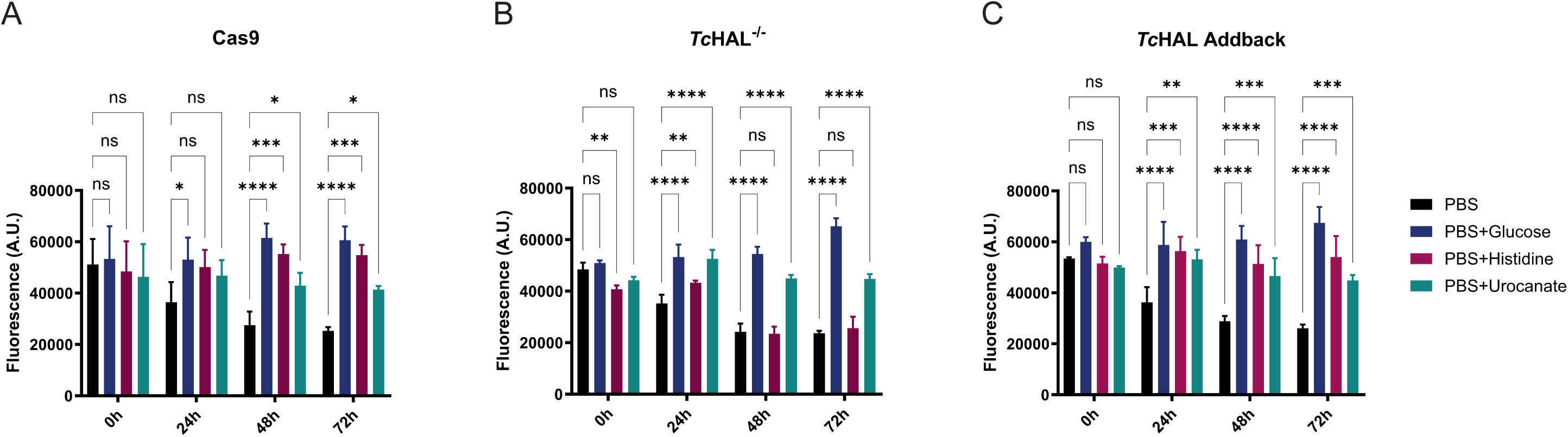
*Tc*HAL epimastigotes are unable to use His to maintain cell viability during nutritional stress. Epimastigotes were kept in PBS or PBS supplemented with glucose, histidine or urocanate. Cell viability was assessed using resazurin over 72 hours. Parasites maintained in PBS and *Tc*HAL^-/-^ epimastigotes maintained in PBS + histidine lose viability over time, whereas cells maintained in the presence of a carbon source remain viable during the course of the experiment. Error bars represent the standard deviation of experiments performed with three independent clones for each cell line (**A.** Cas9, **B.** *Tc*HAL^-/-^ and **C.** *Tc*HAL add-back) for each condition. Statistical analysis was performed using two-way ANOVA with multiple comparisons by Tukey’s test (α=0.05; ns = statistically non-significant difference, * p ≤ 0.05, ** p ≤ 0.01, *** p ≤ 0.001, **** p ≤ 0.0001).

### *Tc*HAL^-/-^ epimastigotes are more resistant to nickel sulphate than parental cell lines

His interacts with divalent metals (Yamashita et al., 1990). In organisms capable of *de novo* synthesis, such as archaebacteria, bacteria, and fungi, this biosynthetic capacity is linked to homeostasis of these metals (Malykh et al., 2018). For instance, in *Aspergillus fumigatus*, disruption of the His synthesis pathway reduces resistance to excess of various heavy metals, including iron, copper, and zinc (Dietl et al., 2016). Given that *T. cruzi* epimastigotes are sensitive to certain divalent metals (de Carvalho & de Melo, 2017), we hypothesized that disruption of the His degradation pathway (through the deletion of the *Tc*HAL coding sequence) might lead to intracellular His accumulation and, therefore, increased metal resistance. To test this, the sensitivity of Cas9, *Tc*HAL^-/-^ and *Tc*HAL add-back strains to nickel was assessed. *Tc*HAL^-/-^ parasites exhibited greater resistance to NiSO_4_ (logIC_50_: 2.688) compared to the Cas9 (logIC_50_: 4.010) and *Tc*HAL add-back (logIC_50_: 2.698) lineages (Supp Figure S2 A-D). These results suggest that the loss of *Tc*HAL expression may lead to intracellular His accumulation, potentially contributing to increased metal resistance.

### Knocking out *Tc*HAL affects the infection capability of the cell-derived trypomastigotes but not of the metacyclic trypomastigotes

Given that *Tc*HAL*^-/-^* parasites are unable to metabolize His, we investigated the impact of disrupting this pathway on the parasités life cycle. First, we assessed the proliferation of epimastigotes over 216 hours. No significant differences in proliferation rates were observed among the *Tc*HAL^-/-^, add-back and Cas9 lineages, indicating that the deletion of *Tc*HAL does not impair epimastigote proliferation under culture conditions (Figure 7A).

**Figure 7.**
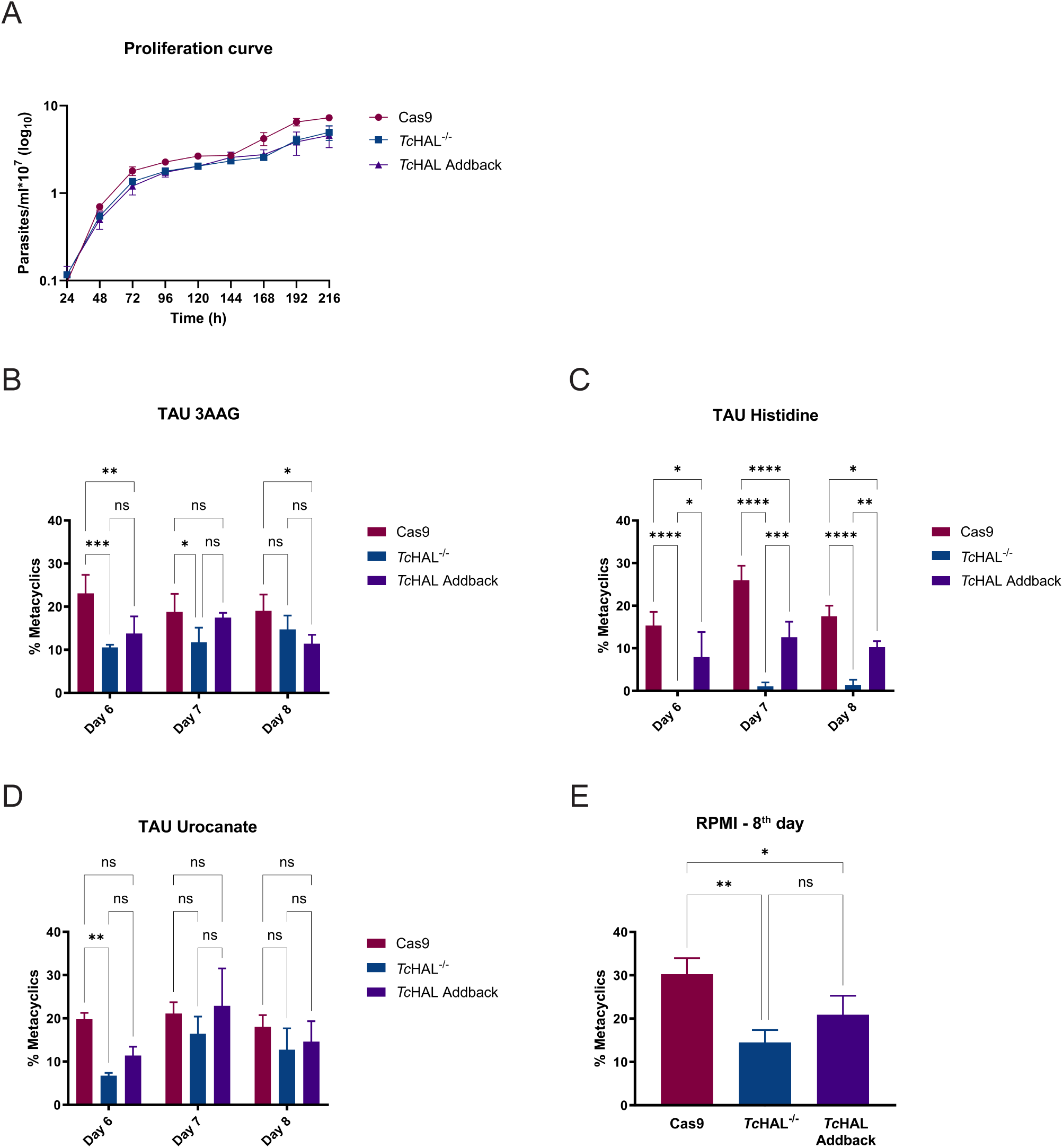
The deletion of *Tc*HAL does not affect the proliferation of epimastigotes but reduces metacyclogenesis *in vitro*. **A**. Proliferation of epimastigotes was monitored by spectrophotometric reading of optical density (620 nm) every 24 hours for 216 hours. When compared to the parental cell line, epimastigotes of the *Tc*HAL^-/-^ and *Tc*HAL add-back cell lines do not show significant differences in proliferation. Error bars represent the standard deviation of experiments performed with three independent clones for each cell line. For metacyclogenesis in TAU, epimastigotes aged in culture for 7 days were subjected to nutritional stress for two hours and then transferred to the desired TAU. Differentiation was monitored by counting in a Neubauer chamber from the 6th to the 8th day after transfer to TAU 3AAG, Histidine, or Urocanate. For differentiation in RPMI, late exponential phase parasites were transferred to RPMI without FBS, at pH 8.2, and kept at 28°C and 5% CO□ for 8 days, at which point the metacyclic count was performed. Knockout lineage parasites have reduced differentiation capacity in **A.** TAU 3AAG, **C.** TAU Uro, and **D.** RPMI, and are completely unable to differentiate in **B**. TAU His. Error bars represent the standard deviation of experiments performed with three independent clones for each cell line (Cas9, *Tc*HAL^-/-^ and *Tc*HAL add-back (Ab)). Statistical analysis of data in graph A, B and C was performed using two-way ANOVA with multiple comparisons by Tukey’s test, α=0.05. Statistical analysis of data in graph D was performed using one-way ANOVA with multiple comparisons by Tukey’s test, α=0.05. (ns = statistically non-significant difference, * p ≤ 0.05, ** p ≤ 0.01, *** p ≤ 0.001, **** p ≤ 0.0001).

Within the triatomine vector insect, *T. cruzi* differentiates from the replicative epimastigote form to the non-replicative but highly infective metacyclic trypomastigote form, a process known as metacyclogenesis. This process can be replicated *in vitro* under chemically defined conditions using a medium called TAU (triatomine artificial urine), which mimics the insect vector environment. TAU is supplemented with various energy sources that support cellular differentiation. The most commonly used formulation, TAU 3AAG, contains three amino acids – Pro, aspartate (Asp), and Glu – and glucose (Contreras et al., 1985). Other energy sources, such as alanine and glutamine, has also been shown to efficient support metacyclogenesis (Damasceno et al., 2018; Homsy et al., 1989). To assess differentiation capacity of *Tc*HAL^-/-^ parasites, *in vitro* metacyclogenesis assays we performed using TAU supplemented with 3AAG, His, or Uro, as well as RPMI medium. *Tc*HAL^-/-^ parasites exhibited reduced differentiation in TAU 3AAG, Uro and RPMI when compared to the Cas9 control, with partial restoration of the phenotypes in the add-back cell line (Figure 7B, D and E). In TAU supplemented with His, the knockout parasites are completely unable to differentiate into metacyclic trypomastigotes, a phenotype that was recovered upon partial restoration of *Tc*HAL (Figure 7C). These results show that the metabolism of His, rather than His *per se*, is essential for the metacyclogenesis observed in His-supplemented TAU.

Since the *Tc*HAL^-/-^ cell line exhibits compromised differentiation capacity *in vitro*, we evaluated its ability to infect the triatomine insect vector *Rhodnius prolixus*. The presence of parasites in the urine, hindgut, and rectal ampulla was assessed. No significant differences were observed in the presence of parasites in the urine of insects infected with the Cas9 and *Tc*HAL^-/-^ cell lines (Supp Figure S3A). Similar results were found for the number of parasites in the posterior gut (Supp Figure S3B) rectal ampulla (Supp Figure S3C) and the percentage of metacyclic forms (Supp Figure S3D). However, when evaluating the number of insects with parasite-negative urine and rectal ampulla, a higher number of insects remained uninfected when exposed to the *Tc*HAL^-/-^ lineage compared to Cas9 control (Supp Figure 3E and 3F).

Finally, given that *Tc*HAL^-/-^ epimastigotes exhibit reduced differentiation capability into metacyclic trypomastigotes *in vitro* across various conditions, we investigated their ability to infect mammalian cells. CHO-K_1_ cells were infected with metacyclic trypomastigotes obtained in RPMI, and the release of cell-derived trypomastigotes (CTs) was monitored from 5 to 15 days post-infection. No significant differences in CT release were observed between the *Tc*HAL^-/-^ and the parental Cas9 cell lines during the initial days of infection (Figure 8A). However, a trend toward reduced release of CT in the *Tc*HAL^-/-^ lineage was noted in the final days of the experiment, with the parental cell line showing higher CT release on days 14 and 15 post-infection. This likely reflects re-infection of CHO-K_1_ cells by CTs. To investigate this hypothesis, CHO-K_1_ cells were infected with CTs from the Cas9, *Tc*HAL^-/-^ or add-back lineages. The release of CTs from cells infected with *Tc*HAL^-/-^ parasites was significantly lower than that of the Cas9 cell line starting from day 10 post-infection, a phenotype that was restored in the add-back cell line (Figure 8B). Interestingly, no differences were observed in the number of infected CHO-K_1_ cells (Figure 8C) or in the number of amastigotes per infected cell (Figure 8D). Together, these results suggest that the reduced CT release in *Tc*HAL^-/-^ infections is not due to defects in host-cell invasion, differentiation, or amastigote replication, but rather arises during the final stages of the intracellular *T. cruzi* cycle.

**Figure 8.**
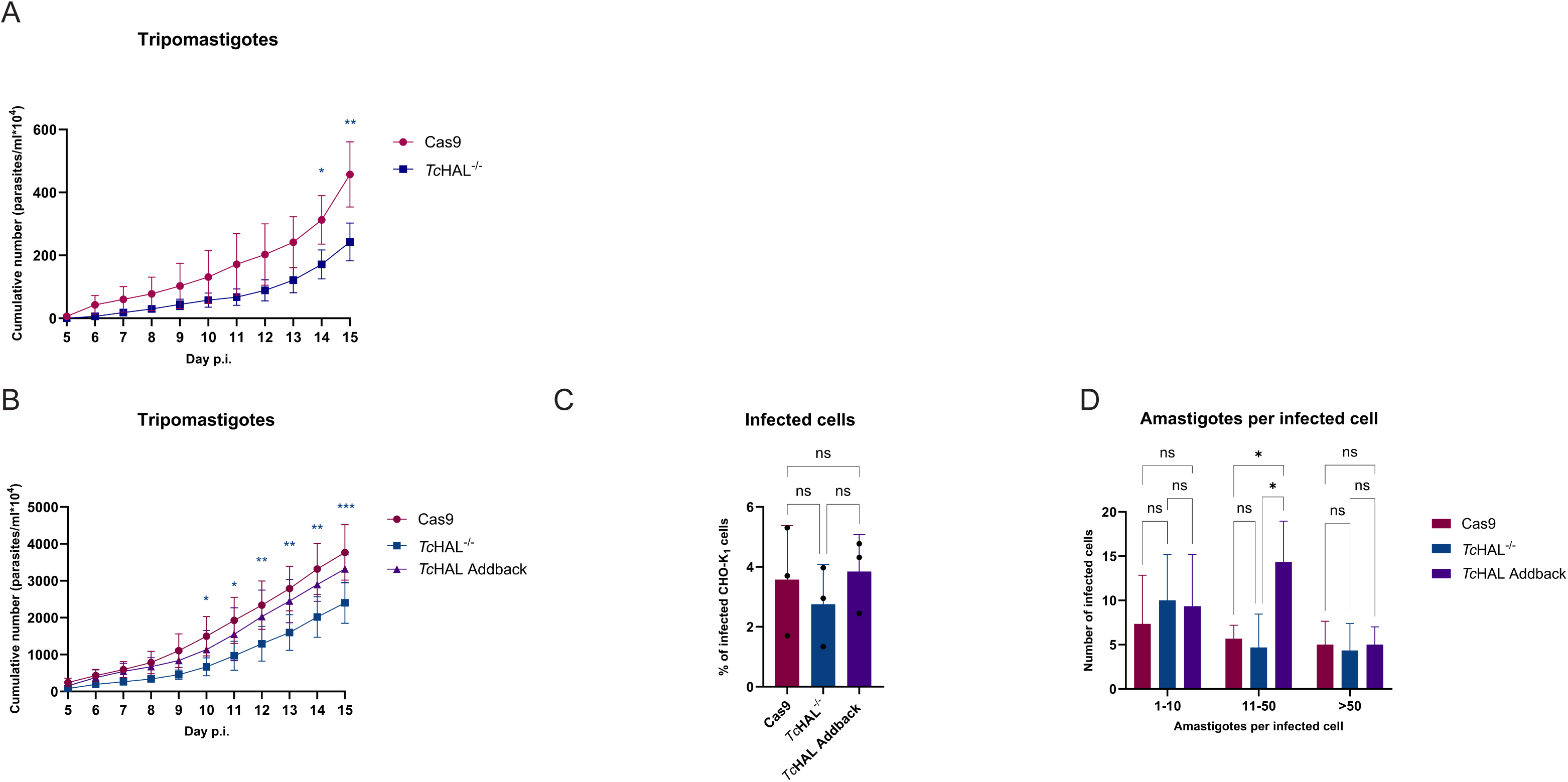
*Tc*HAL knockout affects the infection capability of tissue culture-derived trypomastigotes *in vitro*. **A.** Metacyclic trypomastigotes obtained in RPMI were used to infect CHO-K_1_ cells. Tissue culture-derived trypomastigotes (TCTs) release was monitored by counting in a Neubauer chamber over the period from the 5^th^ to the 15^th^ day post-infection (p.i.). *Tc*HAL^-/-^ metacyclics maintain their ability to infect cells *in vitro* similarly to the control Cas9 up to the 13^th^ day p.i.. **B.** TCTs obtained after a first round of infection with metacyclic trypomastigotes were used to infect CHO-K_1_ cells and TCTs release was monitored. Infections with *Tc*HAL^-/-^ TCTs release significantly less TCTs than Cas9 and *Tc*HAL add-back over time. However, the **C.** number of infected CHO-K_1_ cells and **D.** number of amastigotes per infected cell do not change. Note the difference in the graphs on the y-axis. Error bars represent the standard deviation of experiments performed with three independent clones for each cell line. Statistical analysis of data in graphs A and B was performed using two-way ANOVA with multiple comparisons by Dunnett’s test, α=0.05. Statistical analysis of data in graph C was performed using one-way ANOVA with multiple comparisons by Tukey’s test, α=0.05. Statistical analysis of data in graph D was performed using two-way ANOVA with multiple comparisons by Tukey’s test, α=0.05. (ns = statistically non-significant difference, * p ≤ 0.05, ** p ≤ 0.01, *** p ≤ 0.001).

## Discussion

Amino acids such as His, Pro and glutamine, which play significant roles in *T. cruzi* biology, can be metabolized into Glu, a central metabolic hub. Glu, in turn, serves as a precursor for several key metabolic pathways: (i) it can be converted into α-KG via glutamate dehydrogenase (Barderi et al., 1998; Cazzulo et al., 1977) and (ii) it can undergo transamination by aminotransferases, such as alanine aminotransferase and tyrosine aminotransferase, yielding alanine and α-KG (Vernal et al., 1998; Zelada et al., 1996). The resulting α-KG, as an intermediate of the TCAc, supports ATP production. His in particular, has been shown to stimulates mitochondrial O□ consumption, enhance His-derived CO_2_ production, and maintain cell viability under nutritional stress, indicating an active His-Glu pathway in *T. cruzi* (Barisón et al., 2016). The genes encoding the four-step His degradation pathway (*Tc*HAL, *Tc*UH, *Tc*IP and *Tc*FG) are present in the *T. cruzi* genome. Transcriptome and translatome analysis reveal that these genes are more expressed during the parasités replicative stages (Chávez et al., 2017; Díaz-Viraqué et al., 2023; Smircich et al., 2015). Additionally, structural studies on *Tc*HAL and *Tc*UH revealed that each of them is organized as a homotetramer (Boreiko et al., 2020; Miranda et al., 2020), while *Tc*FG is a homotrimer (Hai et al., 2013).

In most of amino acids degradation pathways, the initial step involves either a transamination or oxidative deamination, yielding the corresponding keto acid as a product. For His catabolism, the first enzymatic reaction involves the non-oxidative elimination of the α-amino group by HAL, producing urocanate and ammonium. HAL enzymes utilize3,5-dihydro-5-methylidene-4H-imidazole-4-one (MIO) as a cofactor, which is synthesised post-translationally through the cyclization of three highly conserved Ala-Ser-Gly (ASG) residues (Louie et al., 2006; Schwede et al., 1999). The crystal structure of *Tc*HAL confirmed the presence of both the ASG tripeptide and the MIO cofactor, demonstrating that the enzyme of *T. cruzi* possesses the essential functional group catalysis, and the post-translational machinery required for MIO synthesis (Miranda et al., 2020). In this study, we determined the kinetic parameters of *Tc*HAL. The measured *K*_m_ and V_max_ values are consistent with those reported for HALs in other organisms (Brand & Harper, 1976; Consevage & Phillips, 1985; Hernandez & Phillips, 1993; Shibatani et al., 1975). Notably, *Tc*HAL exhibits optimal activity in a basic pH range (8 to 10), contrasting with the transporter, which showed higher specific activity at a slightly acidic pH (Barison et al, 2016). This pH preference is consistent with the localization of *Tc*HAL in the acidocalcisomes, where it contributes to the alkalinization of this organelle (Mantilla et al., 2021).

In this study, the generation of *Tc*HAL-null mutants showed that while *Tc*HAL is not essential for the viability of epimastigotes under standard culture conditions, its absence abolishes the ability to oxidize His or utilize it as an energy source to sustain respiration and viability under nutritional stress *in vitro*. Notably, these mutants also exhibited a reduced capacity to incorporate His and/or its metabolic products into macromolecules, indicating that intermediates and end products of His catabolism contribute to anabolic processes in *T. cruzi*. These findings strongly support the idea that the His-Glu pathway is the exclusive catabolic pathway for His in this parasite, supplying metabolic intermediates for ATP synthesis. Interestingly, *Tc*HAL-null mutants retain the ability to transport and utilize Uro, the product of the *Tc*HAL-catalyzed reaction, indicating that downstream steps of the His degradation pathway remain functional. Additionally, *Tc*HAL-null epimastigotes display increased resistance to nickel, a phenotype potentially linked to intracellular accumulation of His.

The process of *in vitro* metacyclogenesis can be induced using TAU, a solution that mimics the urine of the insect vector (Contreras et al., 1985). Supplementing TAU with various substrates has enabled researchers to investigate factors influencing cell differentiation. Amino acids, for example, play diverse roles in this process: some act as “pro-metacyclogenic” (e.g., Pro, Glu, Asp, and glutamine) while others are “anti-metacyclogenic” (e.g., isoleucine, valine, and leucine) when added to TAU (Contreras et al., 1985; Damasceno et al., 2018; Homsy et al., 1989; Nascimento et al., 2024; Rapado et al., 2023). Regarding His, earlier studies reported that the Dm28c strain of *T. cruzi* fails to differentiate in TAU supplemented solely with this amino acid (Contreras et al., 1985). However, our findings reveal that His and Uro support metacyclogenesis in the CL Brener strain, highlighting biological differences among *T. cruzi* lineages. His is abundant in both the hemolymph and excreta of triatomines (Harington, 1961b, 1961a), suggesting it serves as a readily available carbon and energy source for *T. cruzi* stages developing in the insect vector. We further show that *Tc*HAL-null epimastigotes cannot differentiate into metacyclic trypomastigotes *in vitro* when His is the sole oxidizable substrate. Similarly, when Mantilla et al. (2021) mutated the polyP interaction site critical for *Tc*HAL function in acidocalcisomes, epimastigotes failed to differentiate in TAU supplemented with His, ultimately leading to cell death (Mantilla et al., 2021). These results indicate that His acts as an energy and carbon source supporting cell differentiation. However, here we showed that parasite differentiation still occurs in TAU 3AAG, TAU Uro, and RPMI, albeit with reduced efficiency. Additionally, *Tc*HAL deletion does not impair the parasite’s ability to infect and differentiate within the insect vector, where alternative carbon and energy sources are available (Antunes et al., 2013).

Acidocalcisomes, acidic organelles rich in calcium and polyP, play a critical role in ion homeostasis and osmoregulation in *T. cruzi* (Docampo et al., 2011). Under stress conditions, such as hypo-osmotic or alkaline environments, acidocalcisome undergo alkalinization, triggering polyP hydrolysis and calcium release (Rohloff & Docampo, 2006). The polyP content in acidocalcisomes fluctuate during proliferation and across different life stages, suggesting their involvement in the parasite’s life cycle (Ruiz et al., 2001). The localization of *Tc*HAL in acidocalcisomes, along with its polyP-binding activity, suggests that His catabolism plays roles beyond bioenergetics, such as acidocalcisome alkalinization (Mantilla et al., 2021). However, our results show that these functions are not essential for epimastigote survival or differentiation in the insect vector, as the abolishment of *Tc*HAL expression does not significantly impact these processes.

The mechanisms mediating host cell infection by metacyclic and culture-derived trypomastigotes remain incompletely understood (Yoshida & Cortez, 2008). These life stages express stage-specific surface proteins that interact with host cell receptors, triggering diverse intracellular responses in both the parasite and the host-cell (Ferri & Edreira, 2021). Adherence to host cell membranes, an ATP-dependent process, is critical for *T. cruzi* invasion (Martins et al., 2009; Schenkman et al., 1991). One pathway that energetically supports cell invasion is the degradation of L-Pro, which occurs via a two-step enzymatic pathway leading to the formation of L-Glu (Mantilla et al., 2015; Martins et al., 2009), a process homologous to His catabolism. Conversely, L-Pro can be synthesised from L-Glu through a two-step enzymatic pathway (Marchese et al., 2020). Once inside the host cell, trypomastigotes differentiate into replicative amastigotes, escaping the transient parasitophorous vacuole and proliferating in the cytoplasm. After several rounds of replication, these amastigotes differentiate back into trypomastigotes, transitioning through an intermediate stage known as intracellular epimastigote (Almeida-de-Faria et al., 1999). Pro has also been shown to be essential for this differentiation process (Tonelli et al., 2004), however, its transport and intracellular levels are reduced in intracellular epimastigotes (Silber et al., 2002, 2009). This suggests that intracellular epimastigotes may rely on synthesizing Pro from Glu to support cell differentiation. Here, we demonstrate that while His metabolism contributes to *T. cruzi* bioenergetics, *Tc*HAL knockout does not significantly impair metacyclic trypomastigote infection of mammalian cells *in vitro*. Infection with cell-derived trypomastigotes resulted in fewer released trypomastigotes, but no significant differences were observed in the number of infected CHO-K_1_ cells or amastigotes per infected cell. We hypothesize that *Tc*HAL knockout reduces Glu production, thereby compromising Pro synthesis in intracellular epimastigotes, where it is required for differentiation into trypomastigotes. These findings suggest that His metabolism may influence the later stages of *T. cruzi* infection in mammalian cells.

In summary, we showed that *Tc*HAL has kinetic parameters comparable to other HALs in other organisms. Furthermore, the abolishment of *Tc*HAL expression in *T. cruzi* impacts: (i) the ability of epimastigotes to use His as an energy and carbon source; (ii) the process of *in vitro* metacyclogenesis when His is the sole oxidizable substrate; and (iii) the infection competence of cell-derived trypomastigotes. Collectively, these data demonstrate the importance of the metabolism of His in *T. cruzi* for the successful completion of the parasités life cycle.

## Material and Methods

### Cell culture

Epimastigotes of *T. cruzi* CL Brener Cas9 strain (Costa et al., 2018), (a kind gift from Prof. John Kelly, LSHTM) were cultured in liver infusion tryptose (LIT) (5 g L^-1^ liver infusion broth, 5 g L^-1^ tryptose, 4 g L^-1^ NaCl, 0.4 g L^-1^ KCl, 8 g L^-1^ Na_2_HPO_4_, 10 mg L^-1^ hemin, 2 g L^-1^ glucose, pH 7.4) supplemented with 10% heat-inactivated foetal calf serum (FCS), 0.1 mg L^-1^ penicillin, 0.13 mg L^-1^ streptomycin and 200 ug mL^-1^ G418 (Cayman). Chinese hamster ovary cells (CHO-K_1_ cells) were maintained in RPMI medium (Vitrocell) supplemented with FCS 10%, 0.15% (w/v) NaCO_3_, 0.1 mg L^-1^ penicillin and 0.13 mg L^-1^ streptomycin, pH 7.2, at 37 °C and 5% CO_2_.

### Generation of *Tc*HAL null mutants using CRISPR-Cas9 and add-back

Two sgRNA sequences were designed using the “Eukaryotic Pathogen CRISPR guide RNA/DNA Design Tool” (EuPaGDT) (http://grna.ctegd.uga.edu), one to target Cas9 cleavage at the beginning and the other at the end of the *Tc*HAL coding region (TcCLB.506247.220). These sequences were integrated into the forward template primer (Costa et al., 2018), to produce the forward primers sgRNA_Fwd1 and sgRNA_Fwd2 (Supp. Table 1). For sgRNA amplification, a reaction mixture containing 2 μM of each primer (reverse sgRNA scaffold primer and forward primers sgRNA_Fwd1 or sgRNA_Fwd2), 1 × High Fidelity Buffer, 0.2 mM dNTP mix, 2 mM MgSO□, and 1 U of Platinum Taq DNA Polymerase High Fidelity (Invitrogen) was prepared in a 20 μL total volume. The PCR conditions included an initial denaturation at 98 °C for 10 seconds, followed by 35 cycles of 98 °C for 5 seconds, 60 °C for 30 seconds, and 72 °C for 15 seconds, with a final extension at 72 °C for 1 minute.

For the DNA donor amplification, which would replace the gene of interest with the blasticidin resistance gene, primers were designed to include 30 bp homologous sequences flanking *Tc*HAL. The forward primer’s homology sequence began immediately after the start codon of *Tc*HAL, while the reverse primer’s homology sequence began 30 bp before the stop codon (Supp. Table 1). The amplification reaction used 20 ng of plasmid DNA p4432 (de Freitas Nascimento et al., 2018), 0.2 μM of each primer, 1 × High Fidelity Buffer, 0.2 mM dNTP mix, 2 mM MgSO□, and 1 U of Platinum Taq DNA Polymerase High Fidelity (Invitrogen) in a total volume of 40 μL. The PCR conditions involved an initial denaturation at 98 °C for 10 seconds, followed by 40 cycles of 98 °C for 5 seconds, 60 °C for 30 seconds, and 72 °C for 1 minute and 30 seconds, ending with a final extension at 72 °C for 1 minute. Three reactions for each sgRNA and the recombination cassette were pooled, extracted using phenol-chloroform, and precipitated with 0.3 M potassium acetate and 2.5 volumes of isopropanol at −20 °C for 18 hours. Before transfection, the DNA samples were centrifuged at 12,000 x *g* for 30 minutes at 4 °C, washed with 70% ethanol, and centrifuged again at 12,000 x *g* for 5 minutes at 4 °C. The pellet was air-dried in sterile conditions and resuspended in 20 μL of sterile MilliQ water.

To generate add-back cell lines, the pTEX plasmid was modified (Kelly et al., 1992). This plasmid allows for episomal expression of the gene of interest, with selection based on G418 resistance. To replace the resistance gene in pTEX with puromycin, hygromycin, or phleomycin, forward and reverse primers (Supp. Table 1) were designed to re-amplify the pTEX plasmid with a new multiple cloning site, named pTEX_newMCS. The plasmid was amplified by PCR using the Platinum Taq DNA Polymerase High Fidelity (Invitrogen) as previously described, with an extension time of 5 minutes, and then digested with *Nde*I Fast digest (Thermo Fisher Scientific) for 1 hour at 37°C. The plasmid was re-circularized using T4 DNA ligase (Thermo Fisher Scientific), according to the manufacturer’s instructions. Once pTEX_newMCS was confirmed, it was digested with *Bgl*II Fast digest and *Pac*I Fast digest (Thermo Fisher Scientific) for 15 minutes at 37°C, with simultaneous treatment with FastAP Thermosensitive Alkaline Phosphatase (Thermo Fisher Scientific), according to the manufacturer’s instructions. To amplify the fragments corresponding to the puromycin, hygromycin, and phleomycin resistance genes from the plasmids pT-puro (Beneke et al., 2017), pNUS_eBFP_hygro (Tetaud et al., 2002), and p2t7:177 (Alibu et al., 2005), respectively, forward and reverse primers flanked by *Bgl*II and *Pac*I restriction sites were used (Supp. Table 1). The fragments were amplified by PCR and digested with *Bgl*II and *Pac*I for 1 hour at 37°C and ligated to pTEX_newMCS to generate pTEX_puroR, pTEX_hygroR and pTEX_fleoR. The *Tc*HAL coding sequence was amplified using PCR with specific primers flanked by *BamH*I and *Xho*I restriction sites (Supp. Table 1). The amplified product was digested with *BamH*I and *Xho*I Fast digest (Thermo Fisher Scientific) for 5 minutes at 37°C, then ligated to a similarly digested pTEX_puroR plasmid that had been treated with FastAP Thermosensitive Alkaline Phosphatase (Thermo Fisher Scientific). Before transfection, 25 µg of the *Tc*HAL- pTEX_puroR plasmid was precipitated as previously described.

For the transfection, CL Brener Cas9 strain epimastigotes in the exponential proliferation phase were treated with 20 mM hydroxyurea (Sigma) for 18 hours (Olmo et al., 2018). Parasites (4 × 10□) were centrifuged at 1016 x *g* for 5 minutes, washed in 1 mL of transfection buffer (90 mM sodium phosphate, 5 mM potassium chloride, 0.15 mM calcium chloride, and 50 mM HEPES, pH 7.2), and resuspended in 80 µL of the same buffer. This mixture was added to a 0.2 cm cuvette (BioRad) along with 20 µL of DNA for transfection. Nucleofection was performed using the Nucleofector 2b device (Lonza) with a single pulse in program U-033. A control electroporation (without DNA) was done for each antibiotic selection combination. After the pulse, 1 mL of LIT medium supplemented with 20% FCS, 0.1 mg L^-1^ penicillin, and 0.13 mg L^-1^ streptomycin were added to the cuvette. The content was transferred to a flask with 4 mL of LIT medium and incubated for 24 hours at 28°C with 5% CO□. The following day, serial dilutions were performed in a 24-well plate using 60% fresh LIT medium with 20% FCS, 0.1 mg L^-1^ penicillin, 0.13 mg L^-1^ streptomycin, and 40% conditioned LIT medium (previously used to grow epimastigotes for 24 hours in exponential phase) in the presence of the selecting antibiotics (200 µg mL^-1^ G418, 5 µg mL^-1^ blasticidin (Sigma), 5 µg mL^-1^ puromycin (Sigma)). The cultures were maintained at 28°C with 5% CO□ for 3 to 4 weeks. Clones were obtained by limiting dilution, where exponential phase epimastigotes were diluted to a concentration of 2.5 parasites per mL in a mix of 60% fresh LIT medium, 20% FCS, 0.1 mg L^-1^ penicillin, 0.13 mg L^-1^ streptomycin, and 40% conditioned LIT medium, along with the necessary selecting antibiotics. The parasites were then distributed into 96-well plates, with 0.2 mL per well, and kept at 28°C with 5% CO□. After 3 weeks, clones were isolated.

To verify the deletion of the *Tc*HAL coding region, specific primers (Supp. Table 1) were used at a concentration of 0.2 µM, along with 1 × Taq DNA Polymerase Buffer, 0.2 mM dNTPs, 2.5 mM MgCl□, and 2.5 U of in-house-produced Taq DNA polymerase. The amplification conditions included an initial denaturation at 94°C for 3 minutes, followed by 35 cycles of 1 minute at 94°C, 1 minute at 51°C, and 1 minute and 30 seconds at 72°C, with a final extension at 72°C for 5 minutes.

### Preparation of protein extracts

Total protein extracts were prepared from epimastigotes during the exponential proliferation phase. The cells were washed twice by centrifugation at 1,016 x *g* for 5 minutes with PBS 1x (137 mM NaCl; 2.6 mM KCl; 8 mM Na_2_HPO_4_; 1.4 mM KH_2_PO_4_, pH 7.4) and subsequently homogenized in lysis buffer (50 mM Tris; 0.25 M sucrose; 100 mM NaCl; 0.2% Triton X-100, pH 7.6) in the presence of protease inhibitors: 1 mM phenylmethylsulfonyl fluoride (PMSF), 0.5 mM tosyl-L-lysyl-chloromethane hydrochloride (TLCK), and 10 μM N-trans-epoxysuccinyl-L-leucine-4- guanidinobutylamide (E64). The cells were subjected to lysis by sonication in four cycles at 20% power for 30 seconds each, with 30-second intervals on ice between each cycle. The extracts were then centrifuged at 16,250 x *g* for 15 minutes at 4°C. The supernatants containing the protein extracts were quantified using the Bradford method (Hammond & Kruger, 1988), with bovine serum albumin (BSA) solution used as a standard for the calibration curve.

### *Tc*HAL expression, purification and enzymatic activity assays

Expression and purification of *Tc*HAL were performed as described by Miranda et al., 2020. The enzymatic activity measurements of recombinant *Tc*HAL and in total protein extracts were performed as described in Mehler & Tabor, 1953. Briefly, the increase in absorbance at 277 nm, corresponding to the formation of urocanate, was monitored using a SpectraMax I3 fluorimeter (Molecular Devices). The reaction mixture contained: 100 mM Tris-HCl buffer at pH 9, 0.1 mM MnCl_2_, 1.7 mM reduced glutathione (GSH), varying concentrations of His and water to a final volume of 0.2 mL. The reaction was initiated by the addition of each recombinant protein (10 - 50 μg) or total protein extract from *T. cruzi* epimastigotes (50 - 100 μg). The blank for each reaction consisted of the same combination of reagents without adding the purified protein or extract. Absorbance was monitored for 10 minutes at 28 °C, with an initial agitation of 5 seconds. For the kinetic characterization of the recombinant *Tc*HAL the initial reaction velocity (V□) was calculated at short reading times, when the reaction over time can be assumed as near-linear. For obtaining the quantity of Uro formed from the absorbance values, we used the molar extinction coefficient for Uro (CEM_277nm_ = 1.88 M□¹ cm□¹). V□ values were converted into units of product formed per unit of time (μmol/min) and plotted as a function of substrate concentration. The kinetic constants, *K*□ and specific *V*□□□ were determined from the resulting graph, which was fitted to the hyperbolic function described by the Michaelis-Menten model. To assess the pH dependence, the activity of recombinant *Tc*HAL was measured in 100 mM Tris-HCl buffers at a pH range of 6.8 – 9 and in 100 mM CHES buffers (N- cyclohexyl-2-aminoethanesulfonic acid) at pH 9 and 10. The temperature dependence was assessed by varying the temperature of the activity assays between 20 and 55 °C. These data were used to calculate the activation energy.

### CO_2_ trapping

Epimastigotes in the exponential proliferation phase were washed twice by centrifugation at 1,016 x *g* for 5 minutes with 1x PBS, resuspended in 1x PBS, counted, and distributed at a concentration of 2.5 x 10^7^ cells per tube. They were incubated with 10 mM His labelled with 0.1 μCi of L-[^14^C(1,2)]-His for different time points at 28°C. All incubations were performed in biological triplicates with technical duplicates. To capture the CO_2_ released as a product of complete His oxidation, pieces of Whatman paper soaked in 2M KOH were placed at the top of 1.5 mL tubes throughout the incubation period. The papers were subsequently transferred to tubes containing scintillation liquid (Perkin Elmer), and the formation of K_2_^14^CO_3_ on the paper was measured using a Tri-Carb 2910 TR Scintillation Counter (Perkin Elmer). To determine the percentage of labelled His incorporated into proteins, the cells were washed twice with 1x PBS, resuspended in 100 μL of the same buffer, and an equal volume of chilled 1 M perchloric acid was added. The samples were incubated on ice for 30 minutes and then centrifuged for 30 minutes at 16,250 x *g* at 4°C. The supernatant was added to scintillation liquid, and measured by scintillation counting, and the results were attributed to soluble metabolites. The radioactivity incorporated into macromolecules was measured in the pellet resuspended in 0.1% SDS in 15 mM Tris-HCl buffer, pH 7.4.

### Measurement of cell respiration

The respiration rates derived from the degradation of His, Pro (control) and Uro (control) were measured in epimastigotes. Exponential phase epimastigotes were kept in PBS at 28 °C for 16 hours and subsequently recovered for 30 minutes in respiration buffer (MCR: 125 mM sucrose, 65 mM KCl, 10 mM HEPES-NaOH, pH 7.2, 1 mM MgCl_2_, 2 mM K_2_HPO_4_) at 28 °C supplemented or not (negative control) with 5 mM His, Pro, or Uro. Sequential additions of oligomycin A (5 μg mL^-1^) and FCCP (0.3 μM) were used to measure respiratory states 4 and 5. Oxygen consumption rates were measured using intact cells in a high-resolution oxygraph (OROBOROS, Oxygraph-2k, Innsbruck, Austria).

### Assessment of cell viability

The ability of epimastigotes to maintain cell viability under nutritional stress using different energy sources was assessed using the irreversible reduction of resazurin to resorufin (Ansar Ahmed et al., 1994). Exponentially proliferating parasites were washed twice with PBS by centrifugation at 1,016 x *g* for 5 minutes and incubated in the same buffer, either supplemented or not (negative control) with 5 mM glucose, 5 mM His, or 5 mM Uro. Three biological replicates of each cell line were distributed in a 96-well plate at a concentration of 1 x 10^6^ cells per well, with technical triplicates for each condition. Viability was assessed after 24, 48, and 72 hours of incubation at 28 °C by adding 10 μL of a 1% resazurin solution (Sigma). After a 2-hour incubation protected from light, fluorescence emission was measured using a Spectra Max i3 plate fluorimeter (Molecular Devices) with excitation at λ_530_ nm and emission at λ_590_ nm.

### Epimastigote proliferation

Proliferation curves were generated by cultivating epimastigotes in their exponential phase in the LIT medium, as outlined previously, using 96-well plates. The starting concentration was 2.5 × 10□ parasites per mL. Cell proliferation was tracked by measuring the optical density at 620 nm every 24 hours. To evaluate the resistance of epimastigotes to nickel, their proliferation was observed in the LIT medium supplemented with varying concentrations of NiSO□. Each proliferation curve was performed in triplicate (biological replicates), with four technical replicates for each.

### *In vitro* metacyclogenesis

Two protocols were employed to conduct an *in vitro* metacyclogenesis assay. In the first protocol (Contreras et al., 1985), epimastigotes in the exponential proliferation phase were used to initiate a culture with an initial concentration of 5 x 10^6^ parasites ml^-1^, which was maintained in culture as previously described for 6 days. The parasites were washed with PBS 1x at 1,016 x *g* for 5 minutes and then transferred to TAU medium (190 mM NaCl, 17 mM KCl, 2 mM MgCl_2_, 2 mM CaCl_2_, 8 mM potassium phosphate buffer, pH 6) at a concentration of 5 x 10^7^ parasites ml^-1^ and maintained for two hours at 28°C. The parasites were subsequently transferred to one of the following TAU media: TAU 3AAG (TAU + 10 mM glucose, 2 mM aspartic acid, 50 mM glutamic acid, 10 mM Pro); TAU Uro (TAU + 10 mM urocanate), TAU His (TAU + 10 mM His), and maintained for 8 days in a CO_2_ incubator at 28°C. The presence of metacyclics was checked daily by counting in a Neubauer chamber. In the second protocol (Shaw et al., 2016), 1 mL of epimastigote culture in the exponential proliferation phase (5 x 10^7^ cell ml^-1^) was transferred to a culture flask containing 10 mL of RPMI (Vitrocell; prepared according to the manufacturer’s instructions, without the addition of FCS and pH adjustment). The flasks were kept undisturbed at a 30° angle in a CO_2_ incubator at 28°C for 8 days. After 8 days, the number of metacyclics in the supernatant was counted using a Neubauer chamber.

### *In vitro* infection

CHO-K_1_ cells (5 x 10^4^ per well) were cultured in two 24-well plates in RPMI as previously described. After 24 hours, they were infected with metacyclic trypomastigotes or tissue culture trypomastigotes (2.5 x 10^6^ per well). The parasites were allowed to interact with the cells for 3 hours at 37 °C and 5% CO_2_. Following this incubation, the cells were washed twice with 1x PBS to remove non-invading parasites, and RPMI medium supplemented with FCS 10%, 0.15% (w/v) NaCO3, 0.1 mg L^-1^ penicillin and 0.13 mg L^-1^ streptomycin, pH 7.2 was added. The plates were then incubated at 37°C and 5% CO_2_ for 16 hours. Subsequently, the medium was replaced with RPMI containing 2% FBS, and the temperature was adjusted to 33°C. After 72 hours post-infection, the RPMI media was removed from one of the infected plates, and cells were gently washed with 1x PBS. Then, cells were fixed by incubation with a solution of 4% paraformaldehyde (Electron Microscopy Sciences) for 5 minutes at room temperature. After, cells were washed once with 1x PBS and stained with Hoechst 33342 (1:2000 –Thermo Fisher Scientific) for 2 minutes, followed by washing with 1x PBS. Cells were imaged using the fluorescence microscope AMG EVOS XL (Thermo Fischer Scientific). Trypomastigotes present in the supernatant of the second infected plate were counted using a Neubauer chamber, starting from the 5th-day post-infection.

### Triatomine infection assays

The triatomines (*Rhodnius prolixus*) were kept in a controlled environment at 26 ± 1°C, 50 ± 10% RH, and natural lighting. The insects were monthly fed on an artificial feeder heated to 37°C, containing citrated sheep blood obtained from ICTB (Instituto de Ciência e Tecnologia em Biomodelos, Fiocruz, Rio de Janeiro, Brazil). To analyze the infection profile of *T. cruzi* in the intestinal tract of *R. prolixus*, fifth instar nymphs, unfed for 30 days after moulting, were fed using an artificial feeder containing citrated heat-inactivated (56°C/30 min) sheep blood added with a suspension of one of the epimastigote strains of interest (1×10^8^ cell/ml). Twenty days after the infective feeding and moulting to the adult stage, the insects were again fed on uninfected blood. After feeding, the insects were placed in 1.5 ml tubes for two hours for urine collection. The following day, the insects were individually dissected to remove the hindgut and rectal ampulla, which were subsequently homogenized with a pestle in a 1.5 ml tube in 60 μl of PBS to analyze the presence of parasites under a light microscope.

### Graphs and Statistical Analyses

All graphs and statistical analyses were performed using GraphPad Prism v10. The specific statistical tests used in each case are described in the corresponding figure legends.

## Acknowledgements

This work was supported by: Fundação de Amparo à Pesquisa do Estado de São Paulo (FAPESP) 2021/12938-0 (awarded to AMS), Conselho Nacional de Pesquisas Científicas e Tecnológicas (CNPq) grant 307487/2021-0 (awarded to AMS), grant 304862/2022-3 (awarded to AAG) and Wellcome Trust grant 222986/Z/21/Z (awarded to JFN and AMS). We are grateful to John Kelly and Martin Taylor for gently gift of strains and plasmid for the use of CRISPR/Cas9 technology in *T. cruzi*. The funders had no role in study design, data collection and analysis, decision to publish, or preparation of the manuscript. JFN is currently a Wellcome Trust fellow. LM, MJB and JFN were FAPESP fellows and ROOS was CAPES fellow during the development of this work. GTM is currently a FAPESP fellow.

## Supplemental Figure Legends

**Supplemental Figure S1. Expression of recombinant *Tc*HAL.** Fractions from recombinant *Tc*HAL affinity chromatography were resolved on SDS-PAGE. M: molecular mass marker; L: clarified lysate; FT: flow-through; W1 and W2: washes using 60 mM imidazol, E1-5: elutions using 500 mM imidazol.

**Supplemental Figure S2. Epimastigotes of the *Tc*HAL^-/-^ cell line are more resistant to nickel.** The proliferation of epimastigotes was monitored by spectrophotometric reading of optical density (620 nm) every 24 hours for 192 hours in the presence of indicated concentrations of NiSO_4_. Graphs are representative of one of the three biological replicates performed for **A.** Cas9, **B.** *Tc*HAL^-/-^ and **C.** *Tc*HAL add-back epimastigotes. **D.** IC50 was estimated at the 120-hour point. Error bars represent the standard deviation of experiments performed with three independent clones for each cell line.

**Supplemental Figure S3. Knockout of *Tc*HAL does not affect the parasites’ infection capability in the insect vector.** Groups of insects were infected by feeding with epimastigotes of the Cas9 or the *Tc*HAL^-/-^ cell lines. Twenty days after infection, the triatomines were fed again, and urine was collected from the insects for parasite counting (**A**). The next day, the insects were dissected, and the presence and stage of the parasites in the posterior gut (**B**) and rectal ampoule (**C** and **D**) were assessed. The number of insects with urine (**E**) and rectal ampoule (**F**) negative for parasite presence was also evaluated. Note the difference in the y-axis of the graphs. Statistical analysis was performed using unpaired t-tests, using Cas9 as control (ns = statistically non-significant difference).

