## Supplementary figures and images for "Knocking out histidine ammonia-lyase by using CRISPR-Cas9 abolishes histidine role in the bioenergetics and the life cycle of *Trypanosoma cruzi*"

### Supp Figure S1

Supplemental Figure S1

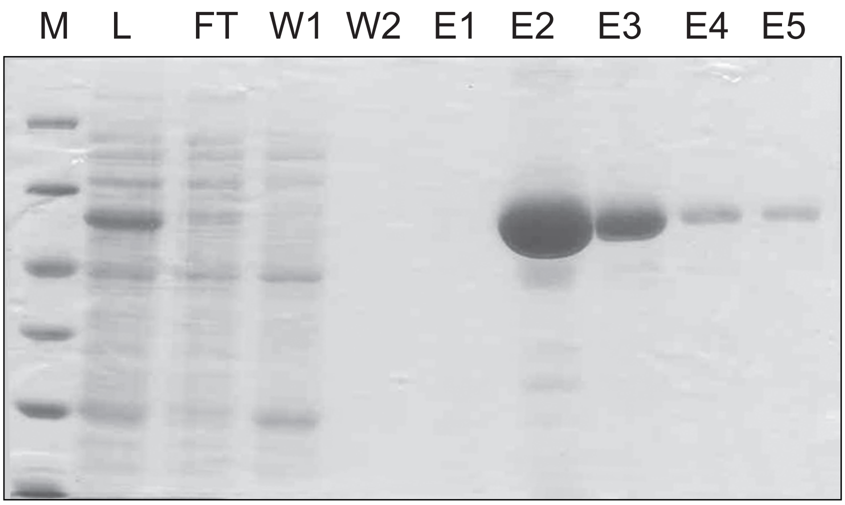

### Supp Figure S2

#### Supplemental Figure S2

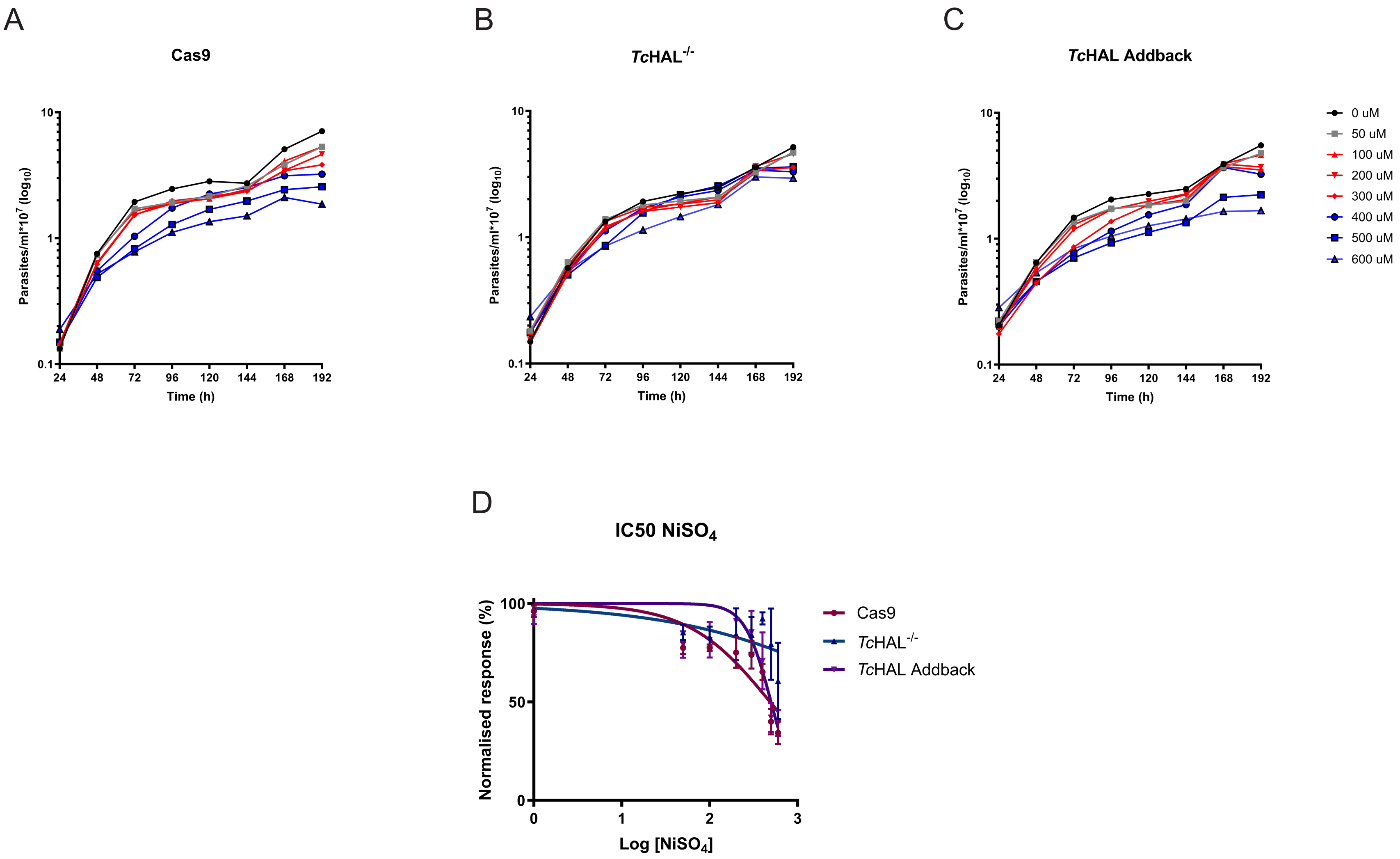

### Supp Figure S3

#### Supplemental Figure S3

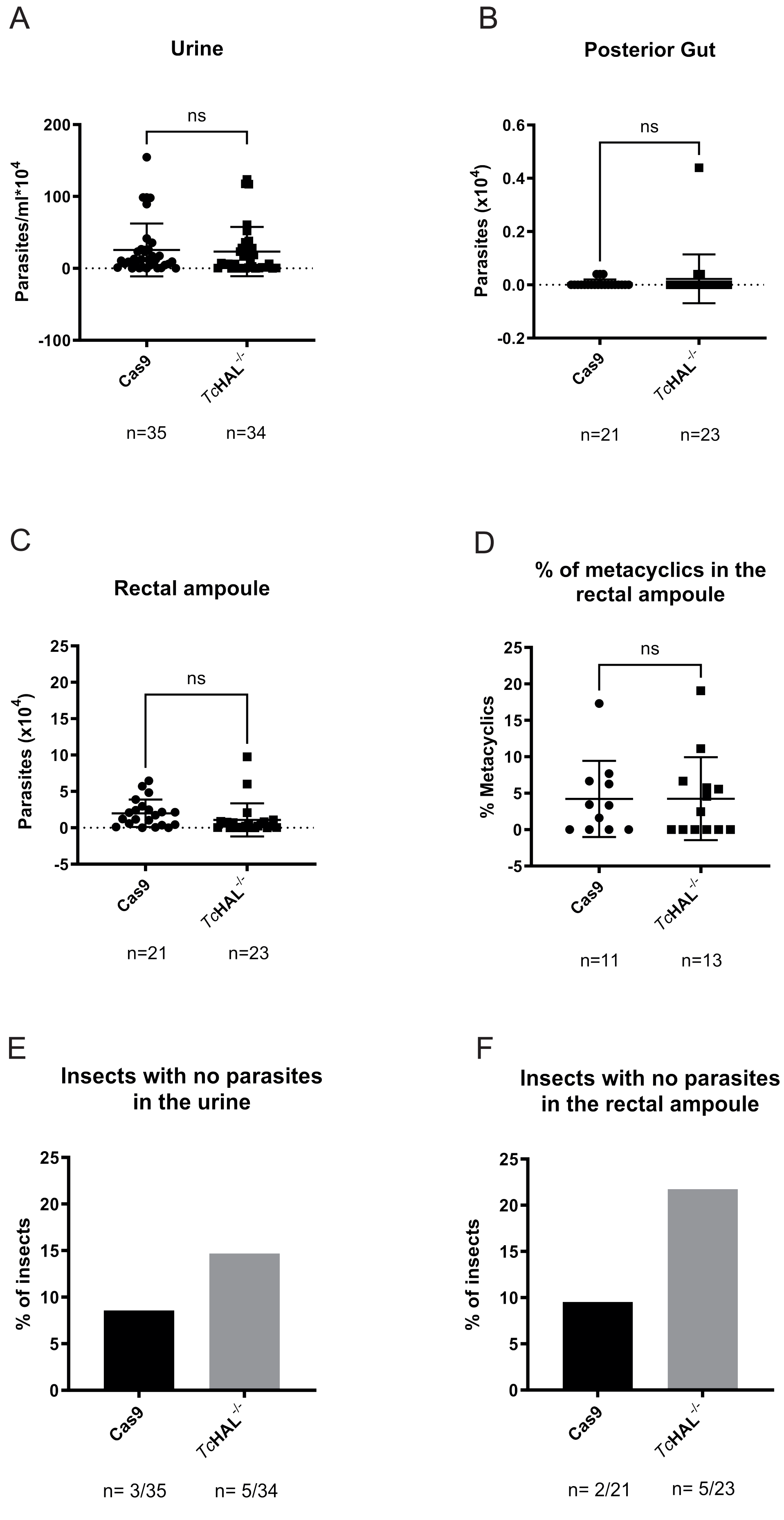
