## Supplementary material for "Knocking out histidine ammonia-lyase by using CRISPR-Cas9 abolishes histidine role in the bioenergetics and the life cycle of *Trypanosoma cruzi*": Supp Table S1

**Table 1. List of primers used for generation of *Tc* HAL null mutants and add-back**

| Primer | Sequence | Description | Use | Tm(°C) |
|---|---|---|---|---|
| sgRNA_Rev | AAAAGCACCGACTCGGTGCCACTTTTTCAAGTTGATAACGGACTAGCCTTATTTTAACTTGCTATTTCTAGCTCTAAAAC | sgRNA scaffold | sgRNA | 69 |
| sgRNA_Fwd1 | GAAATTAATACGACTCACTATAGG*gcccgatgtactgtacgctt* GTTTTAGAGCTAGAAATAGC | sgRNA HAL_33 | sgRNA | 69 |
| sgRNA_Fwd2 | GAAATTAATACGACTCACTATAGGA*ccaagaatcgcgcctcctca* GTTTTAGAGCTAGAAATAGC | sgRNA HAL_1558 | sgRNA | 69 |
| TcHAL_HR_ Fwd | CTGAGGGTTATCCTTGACGGCTGCTCCTTGACGCCATGCCTTTGTCTCAAGAAGAATCCA | Forward primer HAL + blast ORF | Donor DNA | 73 |
| TcHAL_HR_ Rev | TCACATCTTGGATTTCAGCTCAAATGGTTTCTTAACTTAGCCCTCCCACACATAACCAGA | Reverse primer HAL + blast ORF | Donor DNA | 69 |
| TcHAL_Fwd_ *BamH* I | GCAGGATCCATGAGGGTTATCCTT | Forward primer for amplification of TcHAL | Genotyping and cloning | 60 |
| TcHAL_Rev_ *Xho* I | CTCGAGTCACATCTTGGATTTCAGCT | Reverse primer for amplification of TcHAL | Genotyping and cloning | 60 |
| pTEX_Fwd | CATATGTTAATTAAGGGCCCGGGGATCGATCCGTGCACAG | Forward primer for re-amplification of pTEX | Cloning | 70 |
| pTEX_Rev | CATATGGCGGCCGCAGATCTATTGGCTGCAGGGTCGCTCGGTGTTCGAGGC | Reverse primer for re-amplification of pTEX | Cloning | 80 |
| PuroR_Fwd_BglII | AGATCTATGACTGAATACAAGCCAACG | Forward primer for amplification of puromycin resistance gene | Cloning | 57 |
| PuroR_Rev_PacI | TTAATTAATTAGGCTCCCGGCTTACG | Reverse primer for amplification of puromycin resistance gene | Cloning | 58 |
| HygroR_Fwd_BglII | AGATCTATGAAAAAGCCTGAACTCACC | Forward primer for amplification of hygromycin resistance gene | Cloning | 58 |
| HygroR_Rev_PacI | TTAATTAACTATTCCTTTGCCCTCGGAC | Reverse primer for amplification of hygromycin resistance gene | Cloning | 58 |
| PhleoR_Fwd_BglII | AGATCTATGGCCAAGTTGACCAGTGC | Forward primer for amplification of phleomycin resistance gene | Cloning | 62 |
| PhleoR_Rev_BglII | TTAATTAATCAGTCCTGCTCCTCGGC | Reverse primer for amplification of phleomycin resistance gene | Cloning | 60 |
